# Biotic interactions buffer the effects of biodiversity loss on ecosystem functioning

**DOI:** 10.64898/2026.11.20.768031

**Authors:** Blai Vidiella, José M. Montoya

**Affiliations:** Theoretical and Experimental Ecology Station, French National Center for Scientific Research, Moulis, France

## Abstract

Biodiversity loss generally impairs functions on which nature and societies depend upon. Yet, it is difficult to predict under which conditions losing biodiversity has larger effects on functions such as primary production or carbon retention. We present a theory that demonstrates that biotic interactions mediate the intensity of these effects. In species interaction networks with many or strong trophic links, biodiversity loss has weaker effects on functioning because alternative interaction pathways allow surviving species to compensate for species loss. In contrast, in networks with fewer or weaker links, ecosystem functioning is very sensitive to the extinction of species. Here, we identify interaction erosion as a hidden dimension of the biodiversity crisis that provides a unifying explanation for functional consequences of biodiversity loss across ecological contexts.

## Introduction

Biodiversity loss is one of the defining features of our time, with far-reaching consequences for ecosystem stability, resilience, and the provisioning of essential services^1,2^. Evidence accumulated over three decades demonstrates a generally positive relationship between biodiversity and ecosystem functioning (BEF), showing that increasing the number and diversity of species in a community typically enhances functions such as productivity, nutrient cycling, and energy fluxes^3–7^. However, despite the consistency of this broad pattern, the strength and slope of BEF relationships vary markedly across different systems and spatio-temporal scales. This variation is particularly evident in complex ecosystems where multiple interacting trophic levels operate simultaneously, yet the mechanisms underlying this heterogeneity across ecosystems remain incompletely understood^7–9^.

A major reason for this uncertainty in our knowledge is that most BEF research has historically focused on direct links between different aspects of diversity, mostly species richness, and functioning, while paying comparatively limited attention to the role of biotic interactions across trophic levels that characterize natural communities^10–12^. Trophic interactions determine population and community dynamics, regulate biomass fluxes, and dictate the distribution of biomass across levels^13–16^. Biotic interactions can buffer or amplify the consequences of species loss—for example, the removal of a top predator can trigger trophic cascades that disproportionately alter biomass distribution and ecosystem processes^13,17–19^. In addition to their importance for community dynamics and ecosystem processes, biotic interactions are affected by environmental changes^13,16,20,21^. As Janzen expressed over fifty years ago, “what escapes the eye is a much more insidious kind of extinction: the extinction of ecological interactions”^22^. Consequently, if we aim at understanding the effects of the ongoing biodiversity crisis on ecosystem functioning, we need to incorporate the role of biotic interactions.

Much of our current understanding of BEF relationships derives from controlled experiments that simplify food-web complexity, which may not scale-up to real ecosystems that are typically shaped by long-term coevolution and heterogeneous trophic architectures^5,23^. As a result, BEF patterns observed in simplified settings may not reliably capture the dynamics of natural communities, where interaction networks can mediate or even mask the functional consequences of species loss^5,24,25^. This gap between simplified experiments and natural ecosystems becomes critical given that global change is restructuring species interaction networks world-wide^20, 21^.

Most global change drivers—including habitat loss, climate warming, and nutrient enrichment—are altering biotic interactions, ultimately reshaping species interaction networks^12,20,21^. Altered environmental conditions can weaken or rewire predator–prey relationships and shift the relative strength of biotic interactions across trophic levels^18,26,27^. For example, warming often accelerates metabolic rates and alters energy fluxes, while habitat loss reduces the number of realized connections between species, potentially removing the redundant pathways that normally buffer ecosystems against species extinctions^5,9,15^. In some cases, such changes in biotic interactions can amplify the effects of biodiversity loss on ecosystem functioning, leading to steeper BEF relationships. In other cases, shifts on the organization of biotic interactions can trigger a reorganization of remaining interactions that partially compensates for functional loss^19^.

Here, we develop a theory of how changes in biotic interactions within large species-interaction networks shape BEF relationships. Using generalized Lotka–Volterra dynamics in ensembles of random trophic networks (Figure 1), we vary trophic connectance and interaction strength while quantifying total community biomass as the main ecosystem function. This approach highlights how interaction architecture changes the relationship between total biomass and species richness. We predict that highly connected, strongly interacting communities have shallower BEF relationships because alternative interaction pathways permit partial compensation after species loss^8^. Erosion of these interactions should steepen this relationship by increasing the contribution of individual species to total community biomass. We expand the theory to account for additional ecosystem functions, including producer biomass, consumer biomass and trophic transfer.

**Figure 1.**
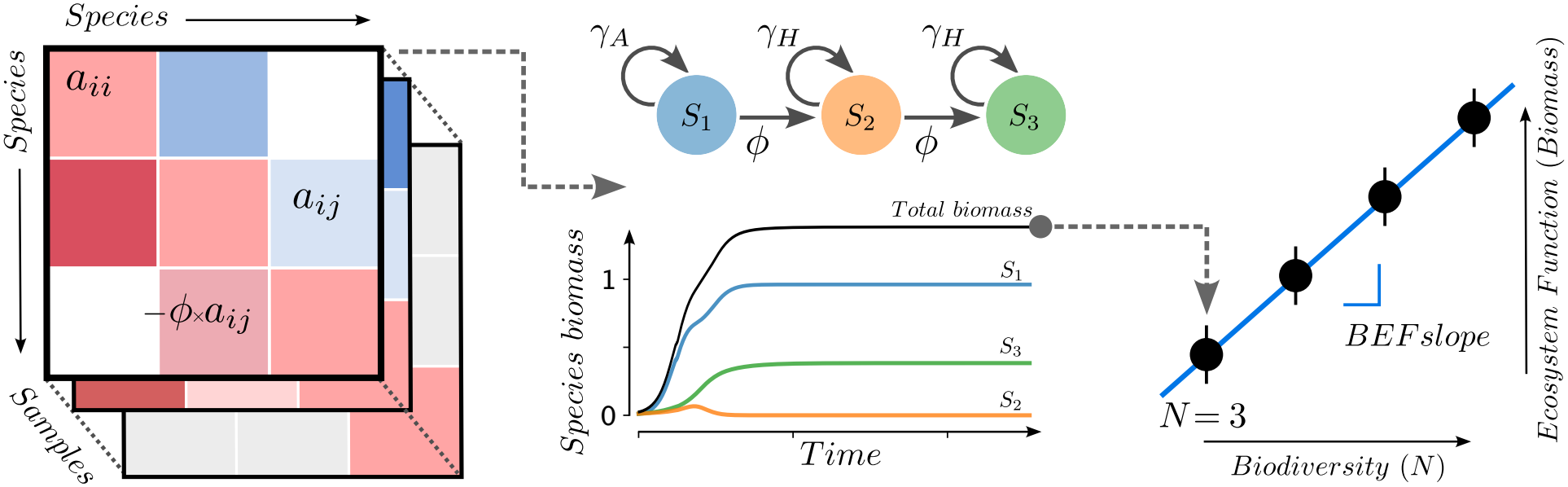
Biodiversity enhances ecosystem functioning through trophic interactions. Conceptual schematic illustrating how increasing biodiversity increases ecosystem functioning. In this manuscript, we measure biodiversity as initial species richness and ecosystem functioning as total biomass (in this example), producer and consumer biomass, as biomass flux. The first panels show multiple model realizations generated from randomly sampled interaction matrices for a fixed parameter set, capturing variability in trophic structure. Each interaction matrix defines a trophic network (centre panel) that governs energy and biomass fluxes. The bottom panel shows a representative Generalized Lotka–Volterra (GLV) simulation for a three-species network, illustrating biomass dynamics through time. Aggregating realizations across richness levels yields the biodiversity–ecosystem functioning (BEF) relationship (last panel).

## Results

### Low connectance amplifies the effects of species loss

Increased network connectance (mean degree, *k*) weakened biodiversity–ecosystem functioning (BEF) relationships, resulting in shallower BEF slopes (Figure 2a). Ecosystem functioning was less affected by species loss in highly connected webs, but declined steeply in sparsely connected networks. Sparser systems, despite this larger sensitivity to species loss, sustained higher total biomass for a given level of diversity. Decreases of BEF slopes with increasing connectance were gradual, without sudden changes.

**Figure 2.**
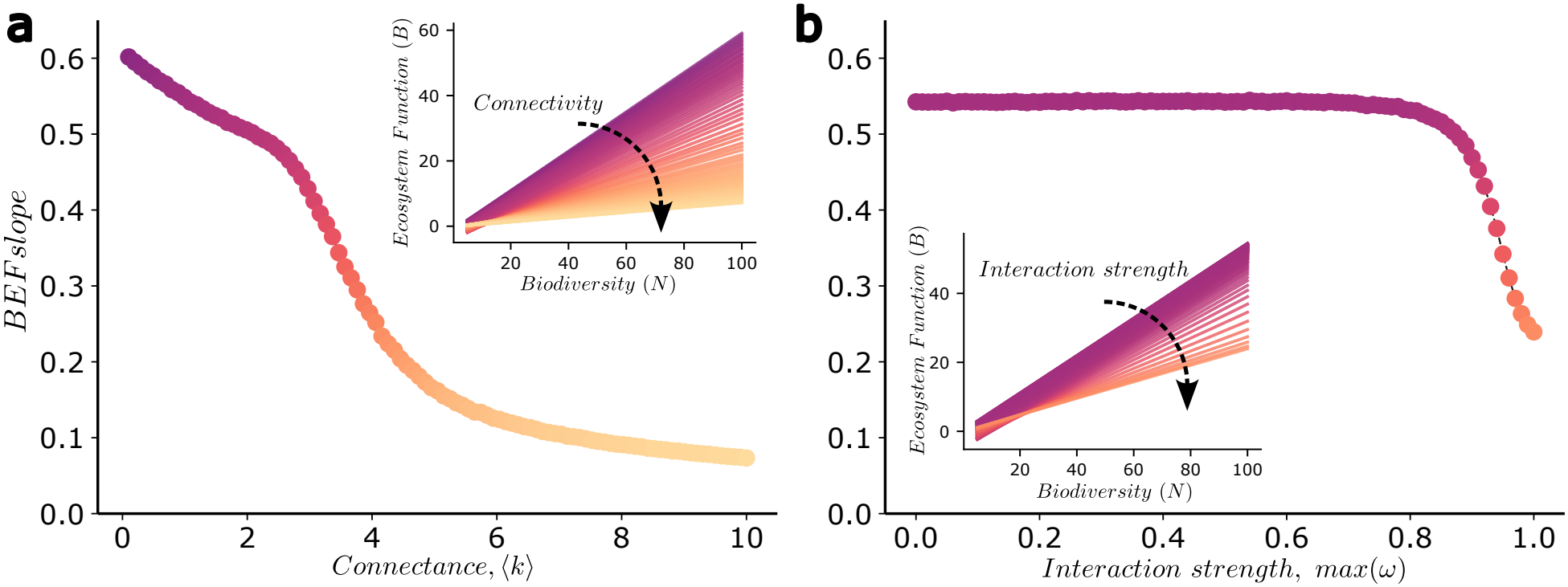
Network connectance and interaction strength buffer ecosystem functioning against biodiversity loss. When network connectance (mean degree, ⟨*k*⟩) is low, biodiversity–ecosystem functioning (BEF) slopes are steep, indicating a strong dependence of ecosystem function on species richness. As connectance increases, BEF slopes become shallower, consistent with greater functional redundancies (panel a). A similar pattern is observed for interaction strength (panel b): weak trophic interactions produce a tight coupling between biodiversity and function, whereas stronger interactions weaken this coupling. For each parameter combination, 1,000 simulations were run across species richness levels. Insets show the corresponding BEF relationships. In panel a, maximum interaction strength was fixed at 0.5; in panel b, mean degree was held constant at 1.

The mechanisms behind this observation lie in how community dynamics change as connectance changes. In highly connected webs, resources are exploited by a larger number of consumers than in sparsely connected webs (see Supplementary Fig. 1), preventing species from reaching their carrying capacities: in the case of resources, due to their higher exploitation by a larger number of consumers, and in the case of consumers, given that they compete with more species for the same resources. Thus, higher consumption and competition in highly connected networks result in smaller biomass values than in sparser networks.

At the same time, these same interaction patterns also shape how ecosystem functioning responds to species loss. Increasing the number of consumer–resource links provides functional insurance against biodiversity loss, as surviving species are more likely to compensate for losses when they interact with more species than when they interact with fewer species^28,29^. This compensatory mechanism can be observed even in small trophic networks, where the change on one species is expected to have important effects on overall functioning. These effects are modulated by network connectance (Figure 3). In poorly connected networks, losing a single species triggers a reduction on total biomass that is much larger than that observed in highly connected networks.

**Figure 3.**
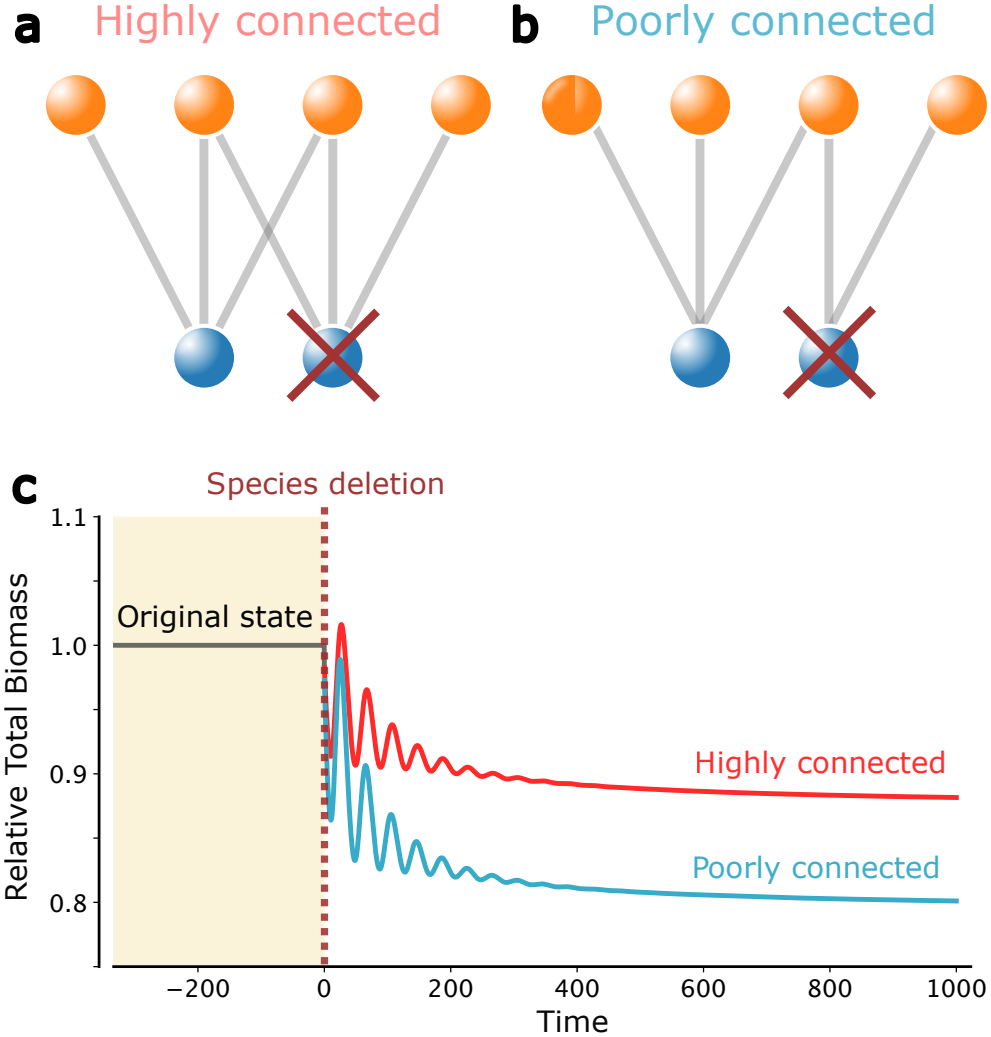
Poorly connected networks respond more strongly to species loss. Network connectance shapes ecosystem responses to species extinctions. Even in these minimal ecosystems, highly connected networks (a) show greater resilience to the loss of a resource species, maintaining higher proportion of ecosystem functioning after the removal of a single species than poorly connected networks (b). As shown in the simulation, both networks respond to the deletion of a resource species (and its interactions), but the highly connected network retains a larger proportion of total ecosystem biomass (c). This difference is consistent with the presence of redundant interaction pathways in more connected networks.

### Weak interactions reduce ecosystem functioning resilience

Weakening interaction strength (*ω*) produced effects comparable to, but distinct from, those of reducing network connectance. Networks with weaker interactions had steeper BEF slopes and higher total community biomass than networks dominated by strong interactions (Figure 2b). This similarity reflects the fact that sufficiently weak interactions contribute little to community dynamics and can functionally resemble the loss of trophic interactions. However, BEF slopes changed less gradually with interaction strength than with connectance. BEF slopes remained relatively high over a broad range of interaction strengths before declining abruptly. This critical value of interaction strength at which this abrupt decline occurred depended on network connectance.

This pattern is consistent with the well-established dependence of community dynamics on interaction strength^30^. As interactions weaken, consumer pressure on resources is relaxed, and producer biomass tends to increase because population dynamics are governed more strongly by intrinsic growth rates and carrying capacities than by consumption^31^. This increase partly reflects a classical predation-release mechanism^32^, whereby reduced consumer pressure allows resources to grow and accumulate. However, predation release alone does not explain the steeper BEF relationships. Weaker interactions are linked to predation release, and they promote producer biomass accumulation (Supplementary Fig. 2), whereas the steeper BEF slopes arise because biomass becomes more dependent on the persistence of individual species. Consequently, networks dominated by weak interactions accumulate more total community biomass^12,14^.

In weakly interacting networks, many species approach the carrying capacities of their resources or of producers, creating weakly coupled components in which biomass accumulates but becomes highly sensitive to species loss. By contrast, in strongly interacting networks, trophic interactions have stronger control over community dynamics, constraining biomass and reducing the contribution of any single species to total community biomass. Thus, weak interactions can increase total biomass while making ecosystem functioning more dependent on the persistence of individual species.

### Combined effects of connectance and interaction strength

We next examined the combined effects of network connectance and interaction strength, asking how movements across this interaction-parameter space influence biodiversity– ecosystem functioning (BEF) relationships (Figure 4). We also compared the effects of losing interactions with those of weakening existing interactions.

**Figure 4.**
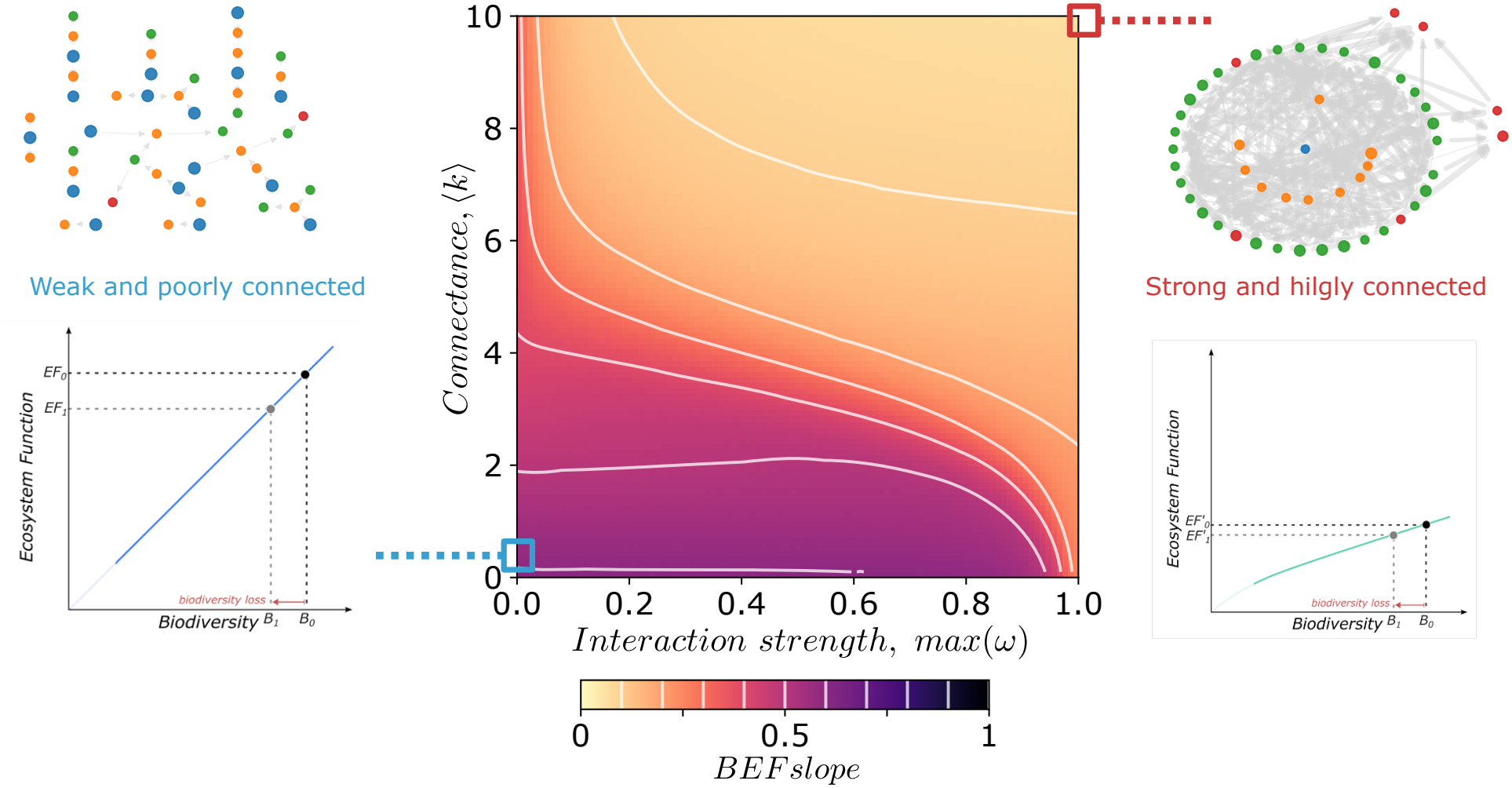
The biodiversity–ecosystem functioning relationship depends on trophic network structure. The biodiversity–ecosystem functioning (BEF) slope declines as biotic interactions become more influential. Increasing either network connectance (mean number of interactions per species) or interaction strength consistently reduces BEF slopes. When biotic interactions are weak (bottom-left region), ecosystem functioning is dominated by autotrophs, and species loss strongly reduces total biomass. In contrast, highly connected systems with strong interactions (top-right region) provide alternative pathways for energy and matter flow, buffering the effects of biodiversity loss. Parameters and simulation procedures are identical to those used in Fig. 2. Results are consistent across a broad parameter space as shown in the Supplementary Figs. 4–6.

Different combinations of connectance and interaction strength produced distinct BEF relationships^19,33^. Highly connected networks with strong interactions showed shallow BEF slopes, indicating that total community biomass was relatively well buffered against species loss. In contrast, sparse networks with weak to moderate interactions showed steeper slopes and greater sensitivity to biodiversity loss. This general pattern was also observed across trophic layers and was robust to changes in both trophic efficiency and growth rates (Supplementary Figs. 4 and 6).

Across most of the parameter space, changes in connectance produced larger changes in BEF slopes than comparable changes in interaction strength (Figure 4). This indicates that losing interactions generally has a stronger effect on the biodiversity–ecosystem functioning relationship than weakening existing interactions. However, in some regions of the interaction-parameter space, relatively small changes in interaction strength produced large changes in BEF slopes. Thus, the relative influence of interaction loss or weakening depends on the original ecosystem configuration, represented here by its position within the interaction-parameter space.

### Biotic interactions and other ecosystem functions

Total community biomass is an integrative and operational measure examining BEF relationships in empirical and theoretical systems^34,35^. However, the same total biomass can reflect very different distributions among trophic levels, from plant-dominated terrestrial systems to consumer-dominated marine food webs with inverted biomass pyramids^36^. We therefore extended our theory to examine producer biomass, consumer biomass and trophic transfer separately. Across gradients of network connectance (⟨*κ*⟩) and maximum interaction strength (max (*ω*)), these analyses test whether interaction structure shapes biodiversity effects beyond the main total-biomass response.

Producer and consumer biomass both depended on network structure (Supplementary Fig. 6), but showed different patterns across the biotic interactions space. Producer-biomass slopes were highest at low connectance, as for total community biomass, whereas consumer-biomass slopes were highest at intermediate connectance and interaction strength. Thus, the interaction-based mechanism extends across trophic properties, while the parameter region of greatest biodiversity sensitivity is property-specific.

We also quantified trophic transfer at the end of the simulations (see Methods). Its BEF slopes closely matched those of consumer biomass: maximum sensitivity occurred at intermediate connectance and interaction strength. This result further illustrates that network architecture channels BEF relationships across trophic properties, although the patterns are not identical^12^.

Finally, we verified that using realized richness (i.e., the number of species persisting at equilibrium) instead of initial richness increases the BEF slopes values but preserves the qualitative patterns across the entire biotic-interaction parameter space. For consistency and comparability with empirical BEF studies, we report results based on initial richness in the main text, while the corresponding analyses using realized richness are provided in the Supplementary Fig. 5.

**Figure 5.**
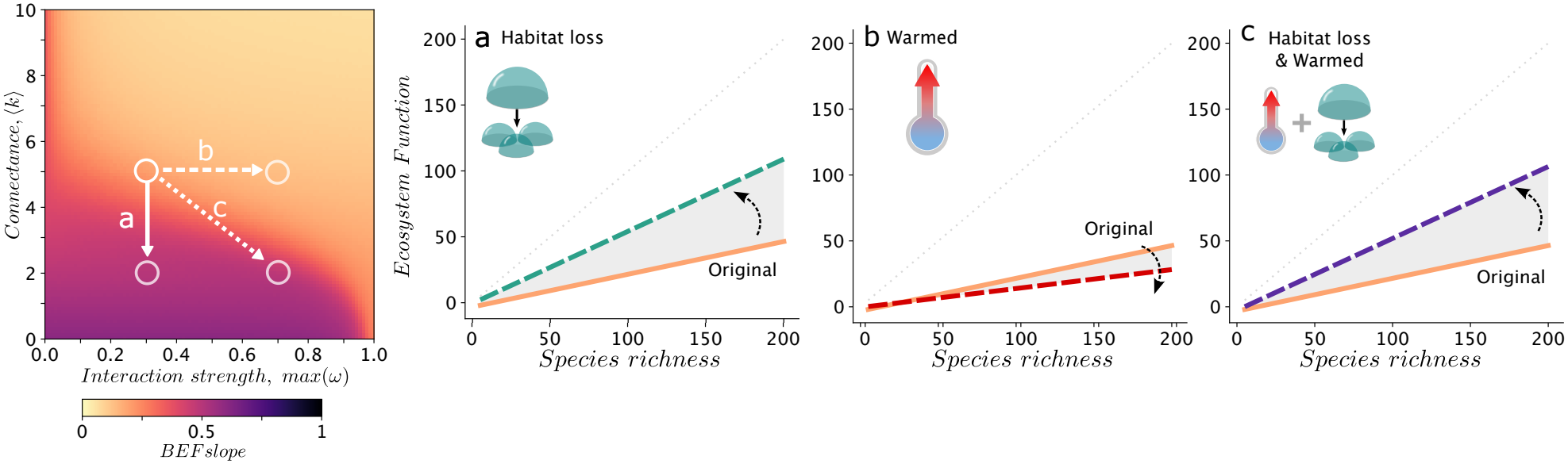
Global change drivers alter the biodiversity–ecosystem functioning relationship. Changes in biotic interactions modify ecosystem resilience to biodiversity loss. Across the biotic interaction (BI) parameter space, variation in BEF slopes reflects changes in functional buffering: shallow slopes indicate weak sensitivity of ecosystem functioning to species loss, whereas steep slopes indicate low resilience. Reduced network connectance, as expected under habitat loss or in small-scale systems, decreases resilience, such that species loss strongly affects ecosystem functioning. In contrast, increased interaction strength, as expected under warming scenarios, reduces overall biomass while constraining species abundances, leading to weaker BEF coupling. The combined effects of these drivers can partially counterbalance one another (see Supplementary Fig. 8).

### Global change effects on BEF relationships

The parameter space of BEF relationships in relation to connectance and interaction strength (Fig. 4) provides a predictive framework for the consequences of global change on ecosystem functioning. Different global change components can shift an ecosystem’s position in this parameter space, altering its BEF relationship. We exemplify this by focusing on the two most pervasive dimensions of global change^20^: habitat loss (and fragmentation) and climatic warming.

Firstly, habitat loss and fragmentation reduces the number of trophic links (Fig. 5a). For example, road density fragments suitable habitats for species and reduces food web connectance^37^. Similarly, recent compilations of empirical networks, including both trophic and mutualistic ones, revealed that a reduction of habitat size reduces both the number of species and the number of links, but the loss of links is faster and more pronounced than the loss of species^38,39^, resulting on reduced connectance with habitat loss. Our framework predicts that reductions in connectance associated to habitat loss and fragmentation will increase BEF relationships, making ecosystem functioning more sensitive to biodiversity loss.

Climatic warming can also modify interaction strength, although the direction and magnitude of this effect depend on the thermal sensitivities of consumers and resources^40,41^. To illustrate one commonly reported scenario, we consider warming that increases interaction strength. In this case, our framework predicts shallower BEF slopes as temperature increases (Figure 5b). This prediction is compatible with empirical evidence that BEF slopes in competitive communities can vary nonlinearly with temperature, including hump-shaped patterns^42^.

Finally, our framework allows us to predict the consequences of global-change components that simultaneously affect the number and strength of interactions, as well as the combined effects of multiple drivers acting independently on connectance and interaction strength. For example, the combined effects of habitat loss and climatic warming increased the BEF slope in our simulations (Figure 5c). This increase was less pronounced than that caused by habitat loss alone because warming acted in the opposite direction by strengthening interactions. The net effect therefore depended on the original ecosystem configuration and on the relative magnitude of the changes in connectance and interaction strength.

## Discussion

Ecosystem functioning relies not just on biodiversity but fundamentally on the network of interactions by which species exchange energy and matter on ecosystems. Our study demonstrates that the erosion of biotic interactions—whether through the loss of connections or the weakening of their intensity—profoundly alters the Biodiversity–Ecosystem Functioning (BEF) relationship. While BEF relationships are hypothesized to vary across spatial and temporal scales^6,7,43^, our theory highlights the critical role of network structure to understand BEF relationships and predict changes due to individual or combined environmental stressors. These findings are consistent with previous theoretical work showing that connectance modulates community robustness to perturbations^44^. While previous studies demonstrated that higher connectance buffers both structural and dynamical responses to species loss, here we extend this insight to ecosystem functioning. We have shown that connectance and interaction strength also constrains the relationship between biodiversity and ecosystem functioning.

If interactions are continually eroded, BEF relationships would progressively increase until they approach a one-to-one relationship. Although this might appear superficially beneficial, suggesting higher functioning for a given number of species, it is not. A tighter relationship actually signals a dangerous loss of functional redundancy, a pattern widely recognized in empirical and theoretical work on ecosystem robustness and interaction loss^33,45–47^. For example, our theory predicts that habitat loss, which reduces connectance faster than species richness^38,39^, increases the BEF slope, making ecosystem functioning more sensitive to further biodiversity loss.

Biotic interactions may help to reconcile the apparent inconsistencies between BEF relationships reported in controlled biodiversity experiments and those measured in natural ecosystems. BEF relationships are typically steeper in experiments^1,5,48^, in part because these studies assemble small species pools with simplified trophic structure and weak dependencies between species-hence, weak interaction strengths-, thereby amplifying species-level effects. In contrast, natural ecosystems are more mature and structurally complex, as they contain richer trophic networks, and a broader and more heterogeneous distribution of interaction strengths characterized by many weak links and a few strong^25,30^. These features tend to dilute the contribution of individual species, resulting in shallower BEF slopes typical from observational studies. Although our model does not aim to fully reconcile these differences, given that most of the experimental settings are plant communities that often involve competitive interactions and artificially high population densities that strengthen interaction strengths, it highlights how variation in network structure can contribute to the contrasting BEF slopes observed across systems.

Our focal result concerns the scaling of total community biomass with species richness. The producer-biomass, consumer-biomass and trophic-transfer analyses show that interaction structure remains consequential across trophic properties, but they also identify a clear boundary condition: the detailed location and magnitude of maximum biodiversity sensitivity depend on the ecosystem property considered. Similarly, we compared the BEF slopes calculated using initial species richness with those based on realized richness, a distinction known to influence BEF patterns in natural systems due to extinction dynamics and species turnover^48,49^. In our case, realized richness produced steeper slopes because species may go extinct but cannot immigrate. Although extinction rates are higher in highly connected and strongly interacting networks, the overall BEF pattern remains the same.

We have exemplified how this framework can inform the consequences of changes on network structure on BEF relationships driven by global change componentshabitat fragmentation and warming in particular. Our examples were simplistic yet informative of the path forward, where more nuanced explorations can expand this framework to spatially explicit models (e.g., cellular automata or partial differential equations), where local interactions and species dispersal jointly drive food web dynamics and functioning. Similarly, more explicit and targeted explorations of the effects of warming on BEF relationships can use species-specific temperature-dependencies on multiple parameters-e.g. growth rates, interaction strengths. Finally, we can increase trophic complexity by embedding habitat- and temperature-dependent effects directly into species interaction, for example incorporating higher-order interactions. Current theory suggests that such mechanisms may introduce additional stabilizing pathways or even amplify destabilizing feedbacks under current global change, offering a promising direction for extending our BEF framework to more precise predictions.

The theory presented here supports and reinforces recent attempts aimed at developing a more holistic, yet practical and operational, perspective for conservation, restoration, and ecosystem management^50^. Under the ongoing biodiversity crisis, it is essential to preserve not only species but the integrity of their interactions^21^ because the erosion of interactions increases the vulnerability of ecosystems for providing and supporting functions. Protecting the density and diversity of biotic interactions—through measures like enhancing habitat connectance, restoring trophic pathways, and conserving behavioral diversity—is essential for sustaining ecosystem functioning in the face of global change^2,11^.

## Methods

### Generalized Lotka–Volterra (GLV) model

We used a standard Generalized Lotka–Volterra (GLV) model to explore the effects of biotic interactions on the biodiversity– ecosystem functioning (BEF) relationship. The GLV framework is widely used to model the dynamics of ecological communities, capturing key features of species interactions and their nonlinear effects on community structure and function^51, 52^.

The GLV model is described by a set of ordinary differential equations governing the growth of each species as a function of its interactions with other species:

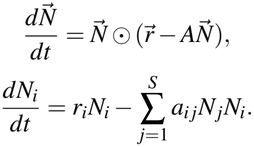

where *N*_*i*_ is the biomass (or abundance) of species *i, r*_*i*_ is its intrinsic growth rate, and *a*_*i j*_ is the coefficient through which species *j* affects species *i*. The code uses the 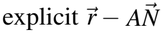 convention above: positive diagonal elements therefore implement self-regulation, with *K*_*i*_ = *r*_*i*_/*a*_*ii*_ in the absence of off-diagonal interactions. For a predator–prey pair, the off-diagonal coefficients have opposite signs, and their relative magnitudes are set by biomass-transfer efficiency.

### Growth rate assignment

Growth rates (*r*_*i*_) were assigned according to trophic position. After constructing the interaction matrix, autotrophs were identified as nodes with zero in-degree. These species were assigned growth rate *γ*_*A*_, whereas heterotrophs received a lower growth rate *γ*_*H*_. Combining this parameter with the self-regulation (*a*_*ii*_) we obtain the upper bound to each species biomass when biotic interactions are not relevant,

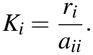

### Maximum interaction strength

Both main axes of our study—interaction strength and connectance—are implemented through the interaction matrix *A*. Self-regulation terms appear on the diagonal and are identical for all species:

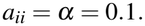

Off-diagonal elements are sampled from a uniform distribution and scaled to set the maximum interaction strength *ω*:

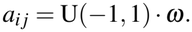

We imposed trophic structure by enforcing reciprocal predator–prey interactions:

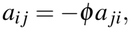

where *ϕ* is a biomass transfer efficiency. Consequently, predator gains are smaller in magnitude than prey losses.

The resulting interaction matrix can be written as

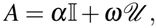

where I is the identity matrix and *U* contains the signed, scaled off-diagonal interaction terms consistent with predator– prey structure.

### Changing connectance

To vary network connectance, we used a Monte Carlo procedure. For each potential interaction pair (*a*_*i j*_, *a* _*ji*_), the probability of retaining the interaction was

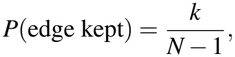

where *k* is the expected mean degree. For each pair, we drew *r* ~ Uniform(0, 1); if ∇ < *k*/(*N* − 1), the interaction was retained, otherwise both entries were set to zero. This procedure generates directed Erdös–Rényi networks with the expected degree distribution and properties described in the Supplementary Note 2.

To avoid isolated species, the sampling procedure was repeated until every node had at least one trophic interaction (in- or out-degree ≥ 1). This ensures that no species is completely disconnected from the network.

Following this procedure, the resulting ensemble spans a wide range of community structures that emerge solely from the stochastic assembly process. As shown in Supplementary Fig. 1, the distribution of species across trophic layers varies systematically with both the mean degree and the total number of species, in agreement with the theoretical expectations derived in the Supplementary Note 2.

### Ecosystem function computation

We quantified two kinds of macro-ecological metrics from GLV simulations: cumulative biomass and biomass flow.

Cumulative biomass captures the long-term aggregate biomass accumulated within the system. We integrated the GLV differential equations until asymptotic dynamics are reached. Then, we computed the time-averaged abundance vector 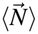 over a long post-transient window to accommodate equilibria, cycles, or chaotic attractors. The total biomass obtained from these persistent abundances, and we partitioned it into:

- **Producers Biomass:** The aggregated biomass of autotrophs or resources (basal species), defined as nodes with an in-degree of zero (*k*_in_ = 0).
- **Consumer Biomass:** The aggregate equilibrium biomass of heterotroph, defined as species that trophically consume at least one other species (*k*_in_ ≥ 1).

Similarly, we computed biomass flow. Biomass flow quantifies the instantaneous energy transfer across trophic links at the final simulation time. We computed the total flux from resources *j* to the consumers *i* as:

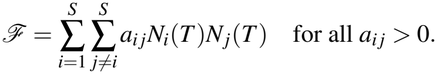

### Biodiversity–ecosystem function slope computation

Once the interaction matrix is generated and the corresponding dynamical equations are defined, we simulate each community until transients vanish and then record species abundances. From the last part of the simulation, we compute mean species biomasses to avoid short-term fluctuations and aggregate them into the ecosystem function of interest (e.g., total, producer, or consumer biomass).

Then, we estimate the BEF slope by fitting a linear regression of ecosystem functioning against biodiversity across all realisations. Where biodiversity is measured either as the initial species richness, *S*, or as the realised richness, defined as the number of species with biomass above 10^−12^ at the end of the simulation. The slope of this fit measures how strongly functioning depends on biodiversity for each network structure.

## Acknowledgements

We thank Alexandre Genin, Renato Moirais, Pauline Narvaez, François Rincon, Jules Treton, Sergi Valverde and the BIOcean5D consortium for discussions. This research was cofunded by the European Union (GA101059915 – BIOcean5D) and by the French Agence Nationale de la Recherche through LabEx TULIP (ANR-10-LABX-41). The views and opinions expressed are those of the authors only and do not necessarily reflect those of the European Union or the granting authority.

## Author contributions

B.V. and J.M.M. conceived the study, developed the theoretical framework, performed the simulations and mathematical analysis, and wrote the manuscript.

